# TACCO: Unified annotation transfer and decomposition of cell identities for single-cell and spatial omics

**DOI:** 10.1101/2022.10.02.508471

**Authors:** Simon Mages, Noa Moriel, Inbal Avraham-Davidi, Evan Murray, Fei Chen, Orit Rozenblatt-Rosen, Johanna Klughammer, Aviv Regev, Mor Nitzan

## Abstract

Rapid advances in single-cell-, spatial-, and multi-omics, allow us to profile cellular ecosystems in tissues at unprecedented resolution, scale, and depth. However, both technical limitations, such as low spatial resolution and biological variations, such as continuous spectra of cell states, often render these data imperfect representations of cellular systems, best captured as continuous mixtures over cells or molecules. Based on this conceptual insight, we build a versatile framework, TACCO (Transfer of Annotations to Cells and their COmbinations) that extends an Optimal Transport-based core by different wrappers or boosters to annotate a wide variety of data. We apply TACCO to identify cell types and states, decipher spatio-molecular tissue structure at the cell and molecular level, and resolve differentiation trajectories. TACCO excels in speed, scalability, and adaptability, while successfully outperforming benchmarks across diverse synthetic and biological datasets. Along with highly optimized visualization and analysis functions, TACCO forms a comprehensive integrated framework for studies of high-dimensional, high-resolution biology.

## INTRODUCTION

Single cell and spatial genomics methods provide high-dimensional measurements of biological systems at unprecedented scale and resolution (Stuart and Satija 2019; Xing et al. 2020). One of the key challenges is attaching meaningful and interpretable representations of the experimental measurements to denote cellular features, such as types, states, cell-cycle stages, or position within tissues (Wagner et al. 2016; Lähnemann et al. 2020) in order to decipher cellular dynamics, communication, and collective behavior. For several biological systems, annotated reference datasets exist that can be leveraged for annotating new, more complex and information-rich datasets.

However, when the new dataset varies substantially from the reference, transferring annotations is a challenging task. A prime example is the transfer of annotations from scRNA-seq to spatial transcriptomics with supra-cellular spatial resolution. For instance, Slide-seq (Rodriques et al. 2019; Stickels et al. 2020) uses spatially scattered beads to measure the combined expression of multiple neighboring cells, possibly of different types. Transferred annotations, like cell type, thus become compositional annotations, such as cell type fractions per bead (**Fig. 1a**). Similarly, for spatial data with sub-cellular or even single-molecule resolution (Zhuang 2021), compositional annotations of local neighborhoods allow segmentation-free annotation and cell assignment of single molecules. Furthermore, compositional annotations can arise not only from cell mixtures but also from ambiguity of categorical annotations (**Fig. 1b**). For example, technical noise, caused by dropout or contamination of ambient RNA can mask the true identity of an individual cell. Finally, biological continua as observed in cell differentiation or regulatory cellular spectra, also require probabilistic cell type assignments (**Fig. 1b**).

**Figure 1:**
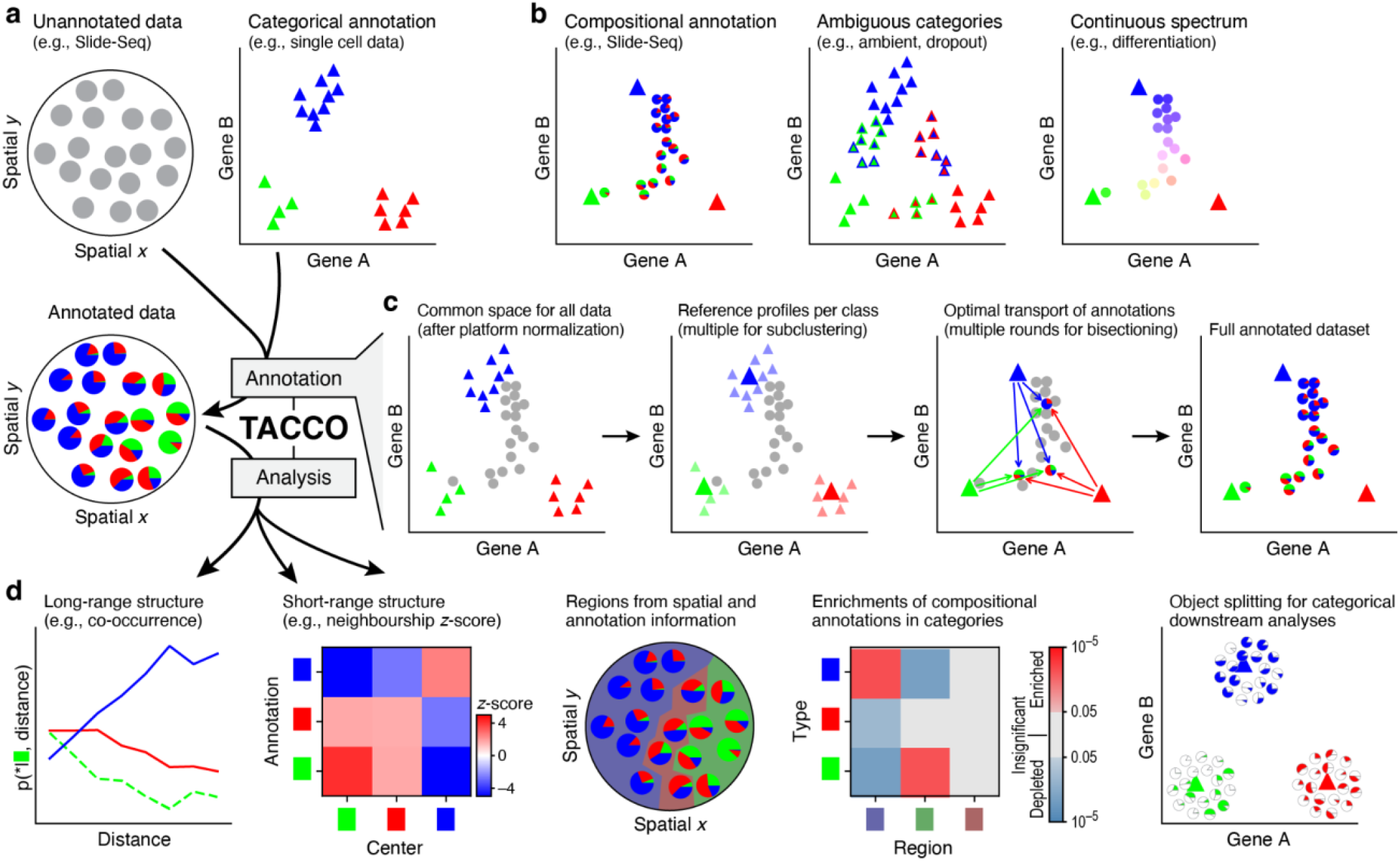
TACCO, a flexible framework for the annotation and analysis of cells and cell-like objects. **(a)** TACCO generates annotations for new datasets of mixtures (top left) using an annotated single-cell reference (top right) and provides methods for downstream analysis of the resulting compositional annotations (bottom left). **(b)** Compositional annotations. Illustrative embeddings of cells and cell-like objects annotated (left) for mixtures (pie charts) of idealized pure contributions (triangles); (middle) as ambiguous annotations (triangles with colored borders) for technical artifacts like high ambient contributions or dropout levels; and (right) continuous annotations (circles) along biological continua. Annotation process. Far left: A labeled reference dataset (*e*.*g*., scRNA-seq data, colored triangles) and a new dataset (e.g., Slide-seq beads, circles) is first presented in a common high dimensional space (e.g., expression space) optionally using platform normalization to make the datasets comparable. Near left: TACCO represents the reference categories by one or multiple representative profiles (large colored triangles). Near right: TACCO uses semi-unbalanced entropic optimal transport to transfer annotations from the reference categories to the new dataset (arrows), generating compositional annotations for the new datapoints (colored piecharts). To improve the capture of subdominant contributions, this process is iterated. Far right: TACCO provides as output compositional annotations for the new dataset. TACCO analysis tools for compositional annotations, especially for spatial data. From left: spatial relationship analysis on long (tissue) and short (cellular neighbourships) length scales; inferring spatial regions by both spatial and annotation information; enrichment of compositional annotations; and splitting compositionally annotated count data into pure contributions for downstream analysis with single-cell analysis tools.

While different decomposition or mapping methods have been developed for specific use cases such as decomposition of spatial measurements (Cable et al. 2021; Achim et al. 2015; Satija et al. 2015; Rodriques et al. 2019; Elosua-Bayes et al. 2021), spatial single-cell mapping (Biancalani et al.; Nitzan et al.; Halpern et al.), or resolving trajectories (Wang and Klein 2021; Schiebinger et al. 2019; Aran et al. 2019; Herman et al. 2018), they are in fact conceptually similar. In particular, in each of these cases, proximity in expression space translates to higher contributions in the annotation, suggesting that it should be possible to develop a general unifying framework across these tasks.

To tackle these challenges, we developed TACCO (Transfer of Annotations to Cells and their COmbinations), a fast and flexible computational decomposition framework (**Fig. 1c**). TACCO takes as input an unannotated dataset consisting of observations (*e*.*g*., the expression profiles of Slide-seq beads) and a corresponding reference dataset with annotations in a reference representation (*e*.*g*., single-cell expression profiles with cell-type annotations), and computes a compositional annotation of the unannotated observations (*e*.*g*., cell-type fractions of the Slide-seq beads). Working in a common high dimensional space (*e*.*g*., gene expression space), TACCO determines these compositions using a variation of the Optimal Transport (OT) algorithm (**Methods**). At its core, OT creates a probabilistic map between the unannotated observations (*e*.*g*., Slide-seq beads) and the means of the classes in the reference representation (*e*.*g*., mean cell-type profiles) according to their relative similarity. Through the OT framework, users can set the similarity metric, constrain or bias the marginals of the mapping, and control the entropy of the annotation distributions (**Methods**). By default, TACCO uses Bhattacharyya coefficients as a similarity metric (**Methods**), which are formally equivalent to the overlaps of probability amplitudes in quantum mechanics, and closely related to expectation values of measurements.

To address concrete biological perturbations and experimental variations between the new and reference data, TACCO provides a set of generic boosters (**Supp. Fig. 1**), including: (**1**) Platform normalization (as in RCTD (Cable et al. 2021)), which introduces scaling factors in the transformation between experimental platforms (*e*.*g*., Drop-seq (Macosko et al. 2015) to Slide-seq (Rodriques et al. 2019)); (**2**) Sub-clustering with multiple-centers to capture within-class heterogeneity; and (**3**) Bisectioning for recursive annotation, assigning only part of the annotation in each step and working with the residual in the next step to increase sensitivity to sub-dominant annotation contributions (**Methods**).

TACCO is also equipped with different analysis tools that leverage the obtained compositional annotations (**Fig. 1d**). It adapts categorical annotation analyses to be applicable to mixture annotations across multiple samples, like the quantification of spatial co-occurrence (Palla et al. 2021) of annotations to analyze long and short range spatial structure, and calculates enrichments of annotations. For spatial data, TACCO can combine spatial and annotation information to define regions with similar annotation compositions across samples. Moreover, TACCO can split compositionally annotated expression data (*e*.*g*., cell-type annotated Slide-seq beads) into categorically annotated expression data (*e*.*g*., split beads per cell-type) using a matrix-scaling algorithm, yielding a data that is amenable to standard downstream single cell analysis workflows.

We evaluated TACCO on four different use cases: (**1**) decomposing cell-type fractions from spatially convoluted gene expression (*in-silico* mixing scRNA-seq profiles for benchmarking, and decomposing colon Slide-seq (Stickels et al. 2020) bead measurements); (**2**) inferring the source cell types of imaged single molecules (without requiring segmentation of cell boundaries from images) for the somatosensory cortex imaged with osmFISH (Codeluppi et al. 2018); (**3**) recovering cell type for scRNA-seq data with harsh dropout and with ambient RNA contamination (on simulated data); and (**4**) predicting differentiation fates of early hematopoietic progenitor cells. As we show below, TACCO consistently achieved performance comparable to or better than other benchmarked methods, and excelled especially in speed and memory consumption.

## RESULTS

We first validated TACCO on simulated Slide-seq data that were generated from an annotated (real) scRNA-seq atlas of the mouse colon as reference (Avraham-Davidi et al. 2022) (**Fig. 2a**) as weighted mixtures of cells drawn from the reference with Gaussian weights parameterized by bead size (**Fig. 2a, Methods**). We measured the *L*2 error between the ground-truth weights and the cell-type fractions inferred by TACCO for varying bead sizes. TACCO matched in reconstruction accuracy with RCTD (Cable et al. 2021), and consistently outperformed all other tested state-of-the-art methods (**Fig. 2a**). Moreover, TACCO was much faster and had lower memory consumption than RCTD, because RCTD fits a detailed model dense in parameters. For example, at bead size 1 (*i*.*e*., bead and cells are of comparable size), the *L*2 errors are 0.081, 0.23, and 0.084 for TACCO, NMFreg (Rodriques et al. 2019), and RCTD, respectively, while TACCO and RCTD runtimes were 84s and 1469s, and memory consumption was 2.7GB and 10.4GB, respectively. Post annotation, TACCO uses the mean reference profiles and the compositional annotation to distribute the actual counts per gene and observation between the cell types and thereby generates separate observations per cell type, keeping the full gene space intact. A subsequent dimensionality reduction then recovers the low-dimensional structure of the reference data (**Fig. 2a, Supp. Fig. 2**).

**Figure 2:**
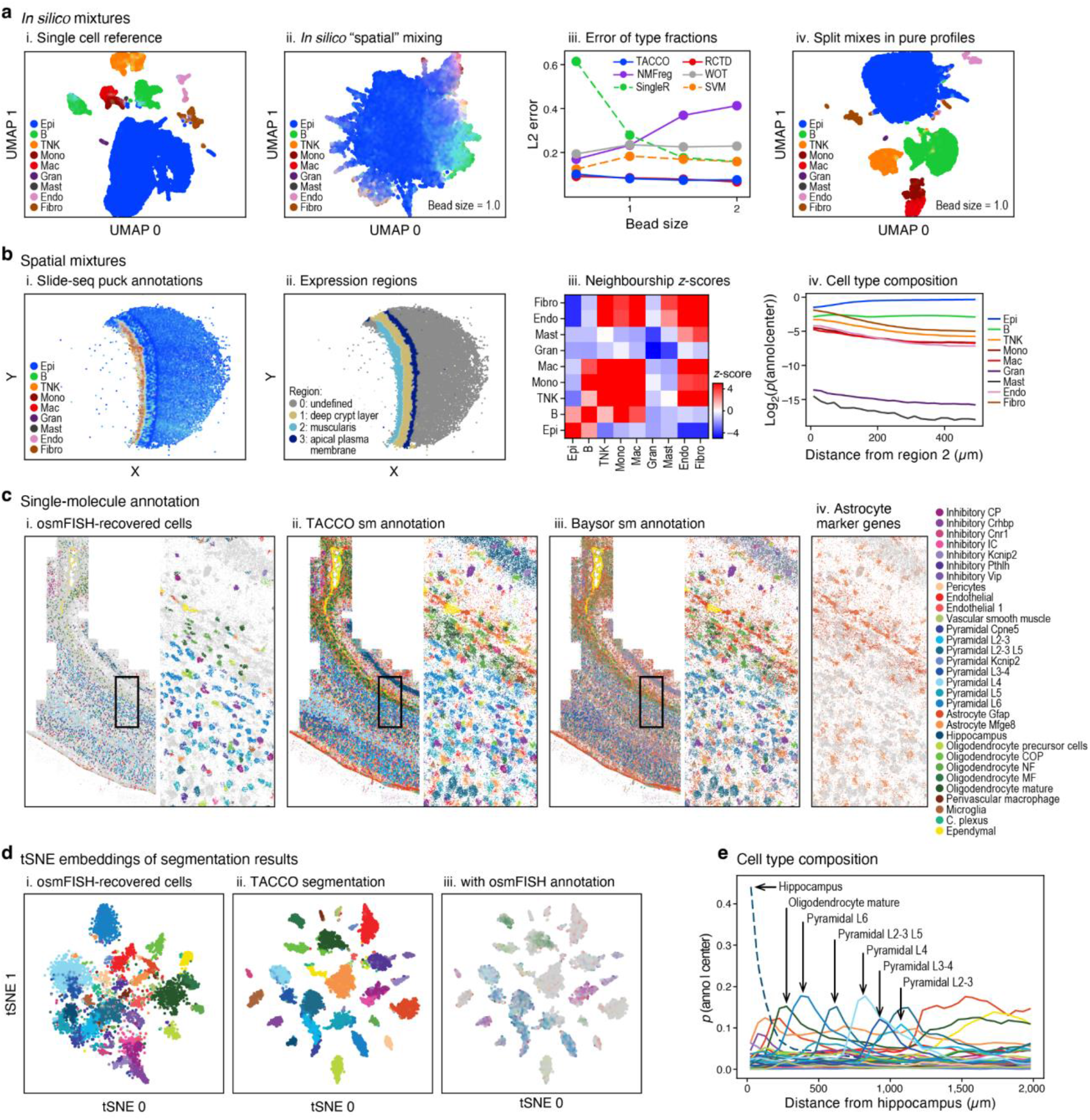
TACCO compositional annotation, cell segmentation and analysis of spatial expression data. **(a)** Analysis of in-silico mixtures. UMAP embedding of scRNA-seq profiles (dots) of mouse colon (i) (Avraham-Davidi et al. 2022), and of in-silico mixed scRNA-seq data before (ii) and after (iv) applying TACCOs splitting procedure into pure contributions. (iii) L2 error (y axis) of cell type annotations for each simulated bead size (x axis). Dashed lines: categorical annotation methods. **(b)** Mapping cell annotations, expression regions and neighborhoods in Slide-seq data with spatial mixtures. (i-ii) a Slide-seq puck of normal mouse colon (Avraham-Davidi et al. 2022) colored by TACCO cell type annotations (i, colors are weighted and summed per bead) or by TACCO-defined regions (ii) (**Methods, Supp. Fig. 3**); (iii) short-range (up to 20µm) neighbourship enrichment z-scores over a background of randomly permuted annotation assignments (color bar) for each pair of cell annotations (rows, columns); (iv) long-range (up to 500µm) dependence of the composition of cell types around beads (log_2_(p(annotations|center)), y axis) at different distances from region 2 (muscularis) (x axis) for each cell type annotation (color). **(c)** Comparison of TACCO and Baysor for single-molecule cell-type-of-origin annotation performance based on single-molecule FISH: (i-iii) Entire section profiled by osmFISH (Codeluppi et al. 2018) (left) and zoom-in of a small region (right) with the positions and cell-type-of-origin annotation of each measured molecule (color, gray molecules are unannotated) using (i) the published annotation of cells from watershed-based segmentation of the poly(A) signal or (ii,iii) the segmentation-free single molecule annotation for cell-type-of-origin by TACCO (ii) or Baysor (iii); (iv) Measured astrocyte marker gene expression on the zoom-in as ground truth for the astrocyte annotation in (i-iii). **(d)** Effective cell segmentation of osmFISH data by TACCO. tSNE embeddings of RNA profiles in cell-like objects from the published watershed-based segmentation (as in c (i)) colored by the published annotation (i), from TACCOs annotation-based segmentation (using the annotations in c (ii)) (ii-iii), colored by either TACCOs single molecule annotations summed per cell (ii) or by the published segmented cell annotation pulled back to single molecules (as in c(i)) and summed per cell (iii) (**Methods, Supp. Fig. 4**). **(e)** Recovery of layered tissue structure in osmFISH data. Long-range (up to 2000µm) dependence of the composition of cell types (p(annotations|center=hippocampus), y axis) recovered by TACCO for TACCO segmented objects at different distances from the hippocampus region annotation (x axis) for each cell type annotation (color).

We also applied TACCO to the annotation of real Slide-seq data from mouse colon with matching scRNA-seq (Avraham-Davidi et al. 2022) (**Fig. 2b, Supp. Fig. 3**). The different characteristics of scRNA-seq and Slide-seq data present annotation challenges. For example, cell composition can vary substantially, due to selection bias (much larger region assayed for scRNA-seq than Slide-seq), or because of differential capture efficiencies (*e*.*g*., lower capture of fibroblasts by scRNA-seq(Slyper et al. 2020) compared to their true tissue prevalence). Moreover, Slide-seq’s low RNA capture rate yields much sparser data than scRNA-seq. Despite these challenges, TACCO recovered the expected layered structure of the colon along the muscularis-apical axis, short-range neighborship relations (closeness of stromal and immune cells with segregating epithelial cells), and long-range gradients on tissue structure scale from muscularis to apical plasma membrane (**Fig. 2b, Supp. Fig. 3**).

**Figure 3:**
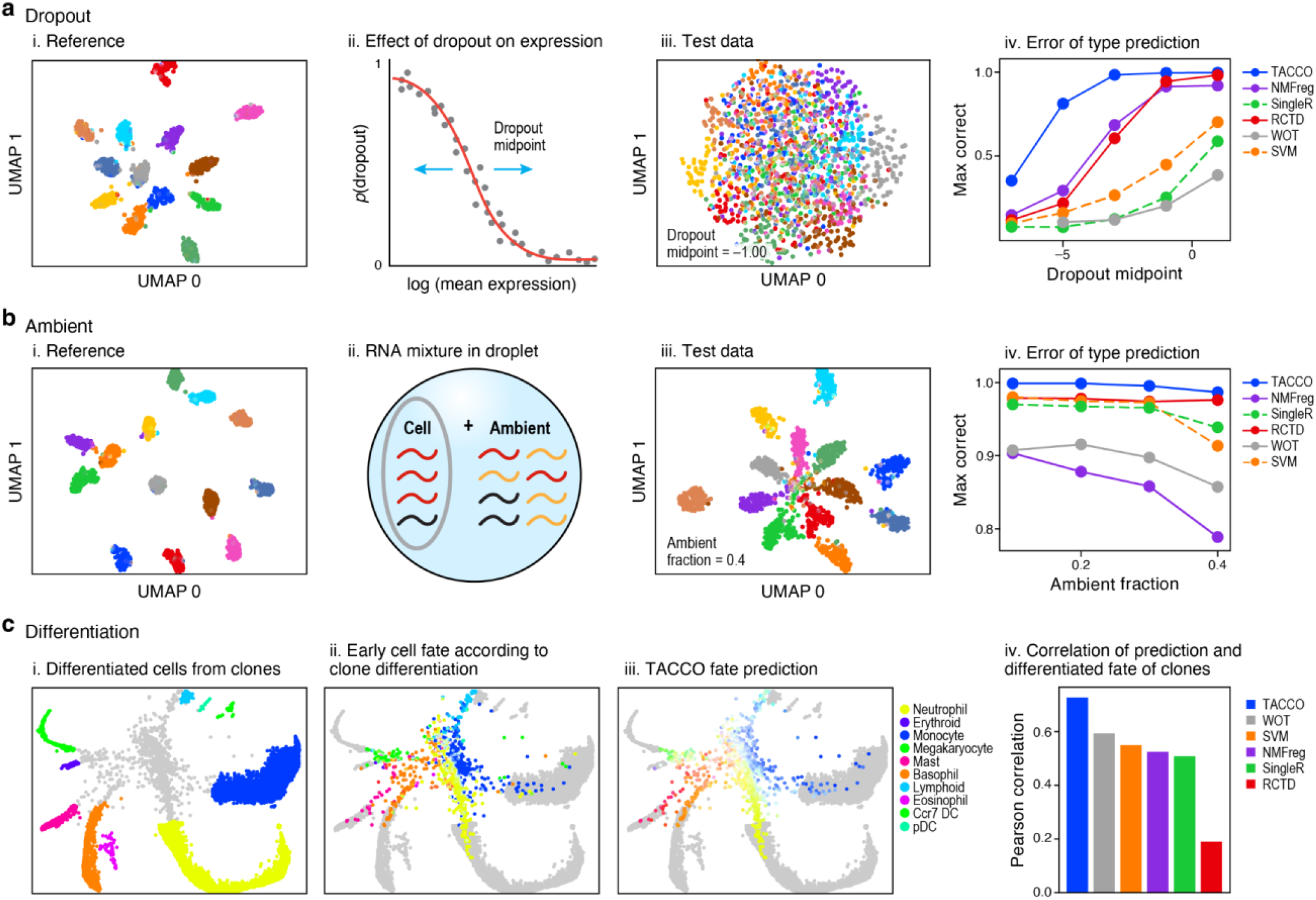
TACCO addresses dropouts, ambient RNA and continuous annotations. **(a)** Dropout task. (i) UMAP embedding of the simulated scRNA-seq profiles without dropout; (ii) schematic of the probability of dropout (y axis) for genes with different mean expression (x axis); (iii) UMAP embedding of the simulated data from (i) but with dropout; (iv) Fraction of correctly annotated cells (y axis, defined by agreement of annotated type with maximal probability and the cell’s true annotation) at different dropout rate (x axis) for different methods (colors). Dashed lines: categorical annotation methods. **(b)** Ambient RNA task: (i) UMAP embedding of simulated scRNA-seq profiles without ambient contribution; (ii) schematic of test data generation, where ambient RNA is added to the cell’s profile; (iii) UMAP embedding of simulated of simulated scRNA-seq profiles from (i) but with additional ambient contribution; (iv) Fraction of correctly annotated cells (y axis, defined by agreement of annotated type with maximal probability and the cell’s true annotation) at different levels of ambient RNA contamination (x axis) for different methods (colors). Dashed lines: categorical annotation methods. **(c)** Differentiation task: (i-iii) Spring plot of scRNA-seq profiles from hematopoiesis(Weinreb et al. 2020) colored by cells types at 4 and 6 day differentiation (i); eventual clonal fate of day 2 cells (ii), TACCO-predicted clonal fate of day 2 cells (iii); (iv) Pearson correlation coefficient (y axis) of predicted and actual fate (as evaluated in (Weinreb et al. 2020)) for each method (colors).

Next, for single-molecule high resolution spatial imaging data (*e*.*g*. MERFISH (Xia et al. 2019) or osmFISH (Codeluppi et al. 2018)), TACCO includes a wrapper for annotating individual, spatially-imaged molecules to cell types, without prior cell segmentation. Standard procedures first segment the captured molecules either based on an image of the cells (Xia et al. 2019; Codeluppi et al. 2018; Eng et al. 2019; Littman et al. 2021) or by finding local maxima in spatially blurred expression fields (Park et al. 2021). Both can be challenging due to high cellular density, irregular cell shapes, and misplaced molecules (Palla et al. 2022; Prabhakaran et al. 2021). TACCO circumvents these limitations by annotating each molecule. To this end, TACCO bins molecules using a Cartesian grid into spatial neighborhoods, computes cell-type annotations for each neighborhood using the reference profiles (as for Slide-seq above), and assigns each molecule an annotation in a manner that recapitulates the neighborhood’s annotation fractions (**Methods**). Applied to a mouse brain osmFISH (Codeluppi et al. 2018) dataset, TACCO successfully accounted for each molecule (**Fig. 2c**), compared to only 36% of molecules annotated following cell segmentation and cell-type classification (Codeluppi et al. 2018) (**Fig. 2c**). Moreover, TACCO matched 59% of the molecular annotations in the original study, compared to only 35% matched by Baysor (Petukhov et al. 2021), a recent single-molecule annotation method based on Bayesian Mixture Models (**Fig. 2c, Supp. Fig. 4**), and was substantially faster (TACCO: 2 mins; Baysor: 19 mins). For cells with non-spheroid shapes, such as astrocytes, single-molecule annotation is especially advantageous: While TACCO and Baysor annotate all molecules, the baseline segmentation and classification approach only accounts for 17% of astrocyte marker molecules (**Fig. 2c**). Subsequent image-free segmentation to cell-like objects based on spatial and annotation information allowed TACCO to recover expression profiles with cell-type annotations, thus utilizing more of the available data and better representing distinct cell types than the baseline as indicated by a higher silhouette score for TACCO than the baseline (0.24 compared to -0.07; **Methods, Fig. 2d, Supp. Fig. 4**). Applying the TACCO segmentation on the Baysor annotation yields an even higher silhouette score (0.45) but at the cost of missing three categories in the annotation (Hippocampus, Inhibitory IC, Inhibitory Pthlh) and a large fraction of molecules in very small segmented objects (28.7% of molecules in objects with less than 20 molecules, compared to 8.3% for TACCO) resulting from a less spatially homogeneous single molecule annotation of Baysor compared to TACCO (**Fig. 2c, Supp. Fig. 4**).

**Figure 4:**
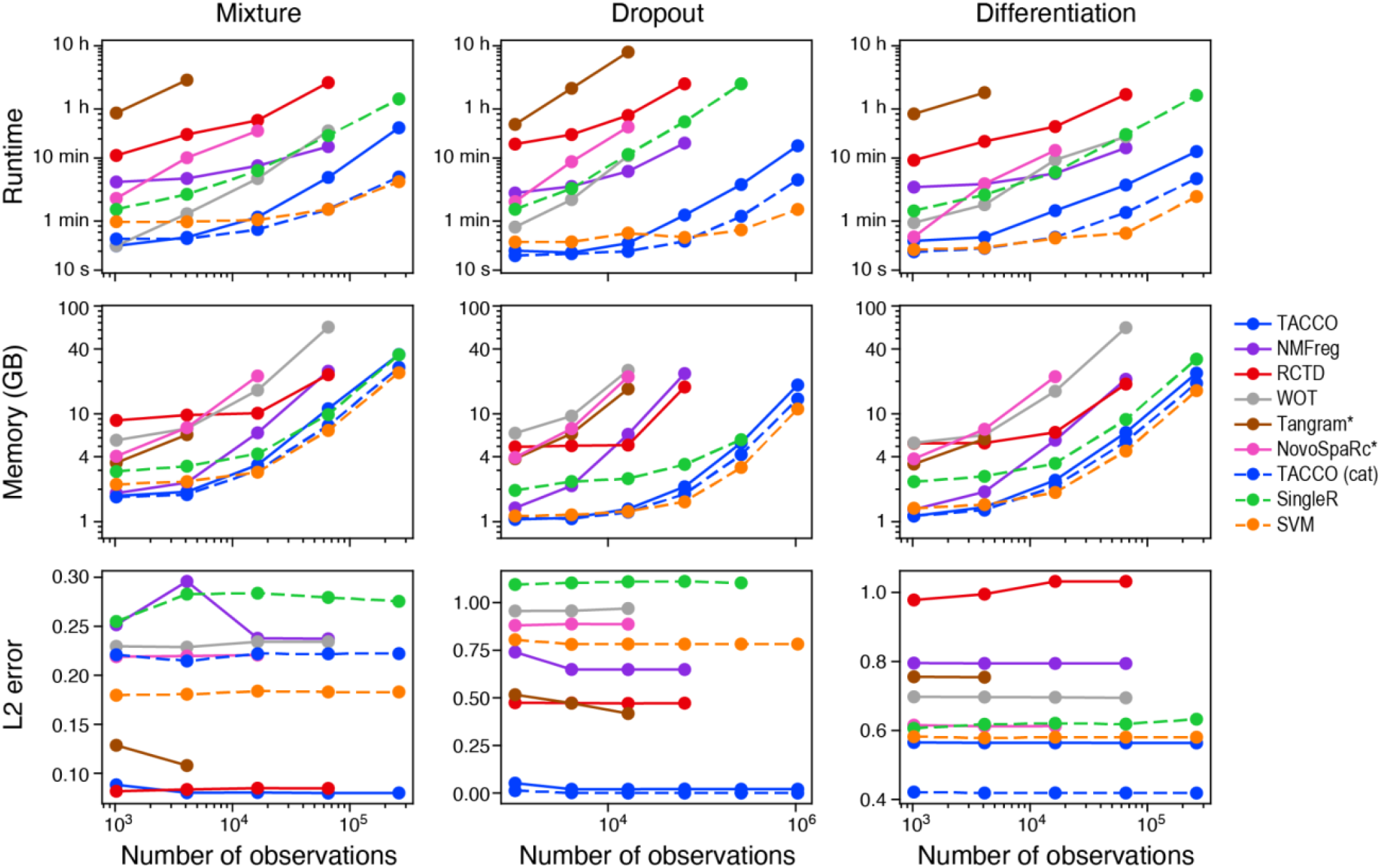
Benchmarking TACCO’s runtime, memory requirement and accuracy of annotation transfer. Runtime (top), Memory (middle) and L2 error (bottom) of each method (colors) in the “Mixture” (left; with beadsize=1.0), “Dropout” (middle; with dropmidpoint=-1.00) and “Differentiation” (right) use cases, at different numbers of observations (cells or beads, x axis), ranging from *2*^10^ ≈ *1k* to *2*^20^ ≈ *1M*, which are up/down sampled from the datasets in Fig. 2. Reference size is fixed at *2*^14^ ≈ *16k*. The maximum compute resources per run are 8 CPU cores for 8 hours with 8 GB memory each. Missing data points indicate that either compute time or memory was insufficient to complete the annotation. Methods with an asterisk (*) do not natively return (fractional) annotations of spatial measurements, which leaves the total annotation fractions in the spatial measurement as degrees of freedom. The wrapper fills that using the reference type fractions. Categorical annotation methods are dashed.

TACCO’s downstream analyses can use this image-free segmented single-molecule dataset to evaluate short- and long-range spatial patterns by computing the cell type composition as a function of the distance to specific annotated cells. For example, we demonstrate this by recovering the layered structure of the mouse brain by calculating the distance to annotated hippocampal cells (**Fig. 2e**). Thus, TACCO’s spatial analyses efficiently bridge spatial scales from single-molecule data to tissue scale.

While TACCO is primarily a compositional annotation algorithm, it can be used as a cell type classifier by interpreting the compositional annotation as probabilities and quoting the class with maximum probability as the classification result. Indeed, TACCO performed well in recovering the ground-truth cell type classification of simulated data with either high dropout rates or high levels of ambient RNAs, as compared to bona-fide classifiers, SingleR (Aran et al. 2019) and SVM, and other compositional annotation methods turned into classifiers (**Fig. 3a**). Specifically, for data with high dropout rates, we simulated a reference dataset of distinct cell types using scsim (Kotliar et al. 2019), where the probability of dropout is a function of log mean expression (Zappia et al. 2017). Although each cell’s type is preserved, increasing this technical noise leads to increasingly “fuzzy” cell-type-specific signals. Based on the fraction of correctly labeled cells, TACCO outperformed both the classification methods and other compositional methods used as classifiers, with larger performance margins in TACCO’s favor for higher dropout rates. In particular, SVM, a top-performing cell-type classification method (Abdelaal et al. 2019), performed poorly when the test data shifted in gene expression. For example, on the same *in silico* dataset with mild dropout rate (dropmidpoint=-1), TACCO correctly assigned 99.9% of cells, whereas SVM only captured 44.7% correctly. For ambient RNA contributions (**Fig. 3b**), we simulated a reference dataset of distinct cell types using scsim (Kotliar et al. 2019) with the ambient RNA model from CellBender(Fleming et al. 2019). TACCO outperformed all other baseline methods, based on the fraction of correctly labeled cells (**Fig. 3b**).

TACCO also resolved continuous biological variability in the context of cell differentiation. To this end, we used a recent dataset (Weinreb et al. 2020) based on the LARRY method, where hematopoietic stem and progenitor cells were clonally barcoded, and clones were followed through subsequent cell division and differentiation, for cells profiled by scRNA-seq at 2, 4 and 6 days. Most early cells (day 2) (96%) were undifferentiated (Weinreb et al. 2020), while day 4 and 6 cells were mostly (61%) differentiated with distinct profiles, thus, for early progenitor cells we construct a proxy of their fate identities based on the distribution of the annotations of their clonal relatives (linked via a shared barcode) (**Fig. 3c**). We then challenged TACCO to predict the fates of early day 2 cells (test data) from the cell-type labeled expression of later, more differentiated cells (reference). TACCO’s predictions are most correlated with the proxy clonal fates (Pearson’s r=0.73), outperforming five other available methods (r=0.19-0.59) (**Fig. 3c**).

We designed TACCO with a focus on practical usability. In addition to its broad applicability to many use cases, TACCO has a modest resource footprint in terms of computing time, memory requirements, and specialized hardware needs. We benchmarked TACCO’s computational requirements relative to those of baseline methods on standard x86 hardware on the “Dropout”, “Mixture”, and “Differentiation” tasks with a range of datasets sizes (10^3^ to 10^6^ observations) (**Fig. 4**). Among all compositional annotation methods in the comparison, TACCO has the lowest runtime and memory requirements, often outperforming the other methods by an order of magnitude or more. Only plain SVM, a categorical annotation method, achieved comparable or better runtime and memory requirements depending on the problem size. However, for categorical annotation, TACCO can also run without the bisectioning booster which improves TACCO’s runtime even further. TACCO achieves all that while maintaining a stable and very competitive L2 error of the annotated compositions with respect to the ground truth across tasks and dataset sizes.

## DISCUSSION

In conclusion, TACCO is a compositional annotation approach and broader analysis package rooted in the realization that multiple tasks in high-dimensional biological representations rely on successfully resolving continuous mixtures over cells or molecules, either due to biological or technical reasons. To leverage annotations of a reference dataset to annotate another dataset, TACCO integrates core, interpretable annotation methods, such as OT, and computational manipulations addressing concrete data perturbations. Together, these yield a top-performing framework in terms of accuracy, scalability, speed, and memory requirements, making TACCO applicable for a wide range of annotation tasks. TACCO could be further extended by modeling continuous reference annotations (e.g., interpolating/extrapolating along a linear reference), by integrating additional priors such as spatial smoothness for annotating spatial transcriptomic data, and by integrating manifold learning techniques to better approximate expression distances from reference profiles. The analytical form of interpretable methods like OT also opens the door to theoretical work on the limitations and guarantees of projecting annotations from one data to another, leading to more informed method selection. While showing here several examples of compositional and categorical annotations, we anticipate that TACCO will help in deciphering many other datasets, such as in decomposing expression of overloaded scRNA-seq experiments (*e*.*g*., MIRACL-seq (Drokhlyansky et al. 2020), which overloads single nuclei in order to capture more rare cells) and in deciphering cells with complex continuous states, such as T cell spectra.

## METHODS

### Overview of TACCO framework

TACCO is based on three guiding principles: modularity, interpretability, and efficiency. TACCOs compositional annotation algorithm is built from a single fast core method, which is then supplemented by a diverse set of wrappers and “boosters”, each providing additional functionality and features. The framework is completed by a set of downstream analysis tools, some of which are especially optimized for analysis involving compositional annotations. TACCO relies on Anndata and seamlessly integrates with the Scanpy (Wolf et al. 2018) ecosystem.

### Compositional annotation

TACCO aims to annotate single cells and cell-like objects, like Slide-seq beads or Visium spots, with compositional annotations. In cases with discrete ground truth (*e*.*g*. a B cell *vs*. a fibroblast) we interpret the compositional annotation as probabilities of belonging to a category. Both objects *b* and categories *t* are in a common high dimensional data space, *e*.*g*. expression space. Generally, objects that are close to a category in that space, should have a high contribution of that category in the compositional annotation.

The goal of the annotation is to find a matrix *ρ*_*tb*_ which gives the distribution of annotation over objects *b* and categories *t*. Some applications imply certain natural choices for the “units” of *ρ*_*tb*_. For example, in the case of count data, the natural units are counts as well, such that *ρ*_*tb*_ gives the counts in object *b* which is attributed to category *t*. The marginal distribution over categories is just the total number of counts Σ_*t*_ *ρ*_*tb*_ = *ρ*_*b*_ per object *b* and is known. The marginal distribution over objects Σ_*b*_ *ρ*_*tb*_ = *ρ*_*t*_ is not known beforehand and is an output of the annotation process. As the marginals per object are known in general, it is equivalent to cite only the normalized distributions 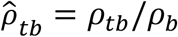 with 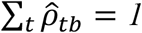 as an annotation result.

### Core annotation method by optimal transport

TACCO’s core method is entropically regularized, partially balanced optimal transport (OT), an intrinsically fast variant of OT). Balanced OT solves the optimization problem *γ*_*tb*_ = *argmin*γ_*tb*_Σ_*tb*_ *γ*_*tb*_*M*_*tb*_ under the positivity constraint *γ*_*tb*_ ≥ *0* and the marginal constraints Σ_*t*_ *γ*_*tb*_ = *c*_*b*_ and Σ_*b*_ *γ*_*tb*_ = *c*_*t*_ with the transport cost Σ_*tb*_ *γ*_*tb*_*M*_*tb*_ =< *γ, M* >_*F*_ given by the Frobenius inner product of a mapping matrix *γ*_*tb*_ and a constant matrix *M*_*tb*_. *M*_*tb*_ encodes the cost of “transporting” or mapping an object *b* to an annotation *t* and must be chosen sensibly to yield a “good” mapping *γ*_*tb*_. The annotation problem is solved if the marginals *c*_*b*_ and *c*_*t*_ and the cost *M*_*tb*_ can be tuned such that *γ*_*tb*_ = *ρ*_*tb*_.

The marginal over annotations is known from the data *c*_*b*_ = *ρ*_*b*_, while *c*_*t*_ = *ρ*_*t*_ is not. This can be used to encode prior knowledge about the data, *e*.*g*. from a reference distribution. To support the general case when such a reference distribution is not available, partially balanced OT is used: instead of fixing the type marginal exactly, a Kullback-Leibler divergence penalty is imposed scaled with a parameter *λ*. This can be used to tune the amount of trust put in the prior distribution.

Choosing a well performing cost function is more ambiguous. The cost function uses the information in data space and assigns a dissimilarity to each object-category combination. A straightforward choice is the cosine distance, or, for expression count data, the cosine distance on normalized and log1p-transformed data. In our benchmarks, the cosine distance on transformed data led to better results (**Supp. Fig. 1**), but the transformation is rather specific for count data. Inspired by the overlap of states in quantum mechanics, a different measure, the Bhattacharyya coefficients, is generally used, with similar performance and without direct reference to counts. Bhattacharyya coefficients are a general measure of the overlap of probability distributions and defined as 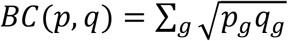 for two probability distributions *p* and *q* in the data space. To allow the user to adapt the method to the needs of their particular application, TACCO implements several other metrics, as well as making all scipy metrics available.

OT’s optimization problem is in general non-convex and numerically expensive. However, using entropic regularization makes it strictly convex and efficiently solvable by using the Sinkhorn-Knopp matrix scaling algorithm. For entropic regularization, the tunable entropy regularization term *ε Ω*(*γ*_*tb*_) = *∈* Σ_*tb*_ *γ*_*tb*_*log*(*γ*_*tb*_) is added to the objective function. This term favors mappings that do not map a given object to a single annotation.

### Alternative annotation methods

The core, optimal transport based annotation method can be swapped by a number of other built-in methods, including non-negative least squares (NNLS) or SVM, or by wrapped external methods from both Python and R, including NMFreg(Rodriques et al. 2019) or RCTD(Cable et al. 2021). Custom core methods can also be added using a functional interface. The wrapped external and custom functions may already include “booster” functionality (below). For example, RCTD already contains platform normalization. Many of the simple built-in methods are amenable to hand optimization, for example by supporting data sparsity consistently, which makes them even faster. All of the built-in methods are generally optimized for standard x86 hardware, but wrapped external methods may use for example GPU acceleration (*e*.*g*., Tangram(Biancalani et al. 2020)).

### Overview of boosters

Boosters can improve the performance of the core method or provide support for ‘missing features’ *e*.*g*., for platform normalization, for using sub-type variability to enhance the type representation in single-cell data, for creating a deconvolution method from a categorical annotation method and the other way round (**Supp. Fig. 1**). They can be combined flexibly to adapt to special requirements for specific applications. The modularity introduced by boosters makes it straightforward to “unbox” TACCO and understand what each part does. While most boosters do have some overhead in runtime, in the end they all do the heavy lifting by calling the fast core method one or several times and transforming its inputs and/or outputs.

### Platform normalization

Dramatic differences in datasets can arise solely from differences in experimental technique that impact the profiled cellular compartments (*e*.*g*., single-cell vs. single-nucleus), cellular compartments (due to differential capture of cells of different types), or gene biases. Annotation of data from one platform using a reference of another platform is much more difficult without accounting for these platform-dependent biases(Cable et al. 2021). This booster can be safely disabled if no platform effects are expected.

A platform normalization step mitigates platform-specific effects by rescaling data from one platform to make it comparable to data from a different platform, separately in each dimension *g* of the data space, *e*.*g*. separately per gene. The necessary rescaling factors are estimated from data. A related but simpler approach compared to (Cable et al. 2021) is pursued, making less assumptions and therefore usable across a broader range of applications.

The representations of the categories *t* as sets of vectors in data space 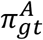 and 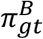 on the two platforms *A* and *B* are linked to each other via the platform normalization factors 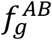 as 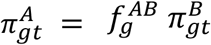. If the category representations 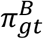 for platform *B* and the (pseudo-bulk) category marginals 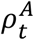 for platform *A* are known, the data space marginals 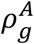 can be written as 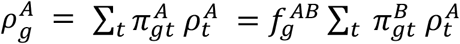, and therefore 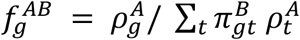. The category marginals 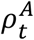 are themselves usually a result of the annotation procedure and used here as input to a preprocessing step for the procedure. However, an iterative scheme can also be used. Starting with the assumption that the annotation marginals are identical to the reference (which is reasonable for example if matched spatial and single cell data are available), most of the gene-wise platform effect can be already captured. Next, platform normalization is re-run after a first round of cell-typing using improved type fractions, and the process is iterated until the normalization factors are stable. In practice, this procedure converges very rapidly with the initial step being by far the most relevant one.

After determining the gene-wise platform normalization factors 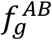, they are used to rescale the data space in either the reference or the test data, or equivalently to work in the units of the test or the reference data. As the probabilities and the resulting annotation distribution are given in terms of these units, choosing test data units or reference data units can lead to quite different results. Which option should be preferred depends on the use case and downstream analyses. Here, test data units are chosen to retain integer count data for object splitting.

### Multicenter

In many cases, there are multiple observations per annotation category in the reference dataset, as is the case in single-cell data. To integrate the variability of these observations while maintaining speed, the observations per category are subclustered with *k*-means clustering to get multiple mean profiles per category. The annotation function is then called with this sub-annotation and its result is summed over the sub-annotations to give an annotation in terms of the original categories. When the reference was already finely annotated, no improvement is expected from this booster, which could then lead to a worse performance, because less data is available per sub-annotation to average noise and the reference gets overfitted.

### Bisectioning

Some annotation methods can be biased towards low entropy annotations, such that they may overweigh dominant categories in mixtures (the extreme case is a categorical classifier). Such methods can be adapted to compositional annotation by running the annotation method iteratively. In each iteration, the annotation is not used as the final result, but instead a certain fraction of it is used to subtract a reconstructed approximation of the data from the original data. The residual is again subjected to the annotation method, while a fraction of the annotation result is added to the final annotation result. This procedure is very similar to gradient boosting. Bisectioning is useful when the objects are additive mixtures of the annotations in the data space. If the data consist (mainly) of objects which are best described by a single annotation, this booster can decrease performance, for example by resolving the ambient contribution in single cell data.

### Object splitting

The annotation method generates an assignment of a composition of annotation categories to every observation. Generally, each observation is associated with more than one category with non-zero contribution. When the categories are cell types, this can be problematic for downstream applications that require the expression profiles of single cells as input, *i*.*e*. pure profiles that can be attributed to a single cell. For example, as cell-type-related expression constitutes a strong signal, analysis of expression programs is easier across cells of shared type. Thus it is desirable to derive several “virtual” observations for every real observation, which correspond to the pure contribution of each annotation category.

A similar idea was proposed in (Cable et al. 2021), where the expected contribution of each cell type to the expression of one Slide-seq bead is approximated. In that approach, cell type fractions that were reconstructed for every bead are taken as input in a Bayesian analysis, along with type-specific expression profiles and the bead-count matrix to yield expected reads per gene, bead, and cell-type. The result, however, lacks consistency with the annotation and the measured count matrix in the sense that the marginal over genes is not recovered. Here, we follow a similar strategy, but we directly integrate marginals as constraints of a matrix equivalence scaling problem in a data-driven frequentist approach.

In this approach, data generation is modeled as drawing single molecules labeled with gene *g*, cell type *t*, and observation/bead *b* from the sample, which constitutes the central object: the joint probability *p*(*gtb*) for a given molecule to be of gene *g*, cell type *t*, and bead *b*. In the actual experiment only *g* and *b* are measured, yielding *p*(*gb*) = Σ_*t*_ *p*(*gtb*), the “t-marginal”. From the reference data, the reference profiles *p*(*g*|*t*) = Σ_*b*_ *p*(*gtb*)/*p*(*t*) are available, and from the annotation process the annotation result *p*(*tb*) = Σ_g_ *p*(*gtb*), the “g-marginal” is obtained.

*p*(*gtb*) is modeled with a Bayesian-inspired product ansatz: *p*(*gtb*) = *p*(*gt*) *n*(*gb*) *n*(*tb*), with free parameters *n*(*gb*) and *n*(*tb*). These are fixed by enforcing the t- and g-marginals.^1^ The ansatz can be interpreted as adjusting the cell-type profiles and pseudo-bulk annotation with object-wise scaling factors per data dimension *g* and annotation category *t*, such that the measurement and the annotations are reproduced exactly. To determine the parameters from the marginals we have to solve a separate matrix equivalence scaling problem per object *b*.

Matrix equivalence scaling problems are guaranteed to have a solution if the matrix to be scaled, *i*.*e. p*(*gt*), has only positive entries. Therefore, a small positive number is added to all the elements of *p*(*gt*) to make the problem well-defined. These problems can be solved by iterative normalization of the columns and rows of *p*(*gt*). This simple algorithm is known under many names, *e*.*g*. the RAS algorithm or for doubly stochastic matrices as the Sinkhorn-Knopp algorithm, and it is also the algorithm used to solve OT efficiently. In contrast to OT, where there is a single matrix scaling problem, here a separate problem is solved for every object. Although this initially seems like a practical performance problem, these problems can be solved in parallel and use the same data, leading to speedups by reducing memory accesses. Moreover, the sparsity of the count frequency matrix *p*(*bg*) can be used to implement the RAS algorithm very efficiently for the problem at hand.

By rescaling the resulting *p*(*gtb*) with the total weight per object (*e*.*g*., total counts per cell for sequencing data) a consistent annotation-resolved split of the measurement is obtained, consisting of floating point numbers. For expression count data, this split generally includes many values much smaller than 1. To optimize sparsity and obtain integer counts, an option is available to round this result by flooring and re-distributing the remaining reads (via multinomial sampling from the remainders). The resulting split count matrix retains biological signal and can be used in standard downstream analyses (**Supp. Fig. 2**).

### Single molecule annotation

To annotate single molecules by cell type assignment, TACCO first bins the single molecule data in space to generate aggregate cell-like objects. TACCO then annotates them either using an internal or wrapped external methods. Subsequently, TACCO maps the resulting compositional annotation back to the single molecules, as follows. First, object splitting is used to distribute molecules to annotations, resulting in a probability for every molecule species in the bin to have a certain annotation. This annotation is then distributed randomly among the molecules of that molecular species, such that each molecule has only a single annotation and the population of molecules reproduces the annotation probability. Because spatial binning can introduce arbitrary boundary artifacts in the definition of local neighborhoods for molecules, TACCO repeats the binning and annotation for *N* (usually 2-3) spatial shifts of the cartesian grid in steps of *1*/*N* of the grid spacing per spatial dimension. For *d* spatial dimensions this results in *N*^d^ annotations per molecule. The final annotation of each molecule is then determined by a majority vote. This simple single-molecule annotation is only feasible with fast annotation methods, which can be run multiple times on differently binned data in a reasonable amount of time.

### Image-free segmentation

An image-free, density-based cell segmentation approach uses the per-molecule annotation to determine molecule assignments at critical cross category boundaries. We assume that the exact assignment of boundaries within a category is not very important as long as it gives rise to a reasonable size of the segmented objects. The segmentation is implemented as graph-based hierarchical spectral clustering.

As the number of single molecules can easily be of the order of *10*^9^, an efficient Euclidean sparse distance matrix computation is required. As scipy’s sparse distance matrix calculation is serial and too slow for this application, a custom, fast, parallel, and still generally applicable algorithm was developed and implemented. After calculating the distances 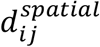 between molecules *i* and *j* in position space, a distance contribution is added in quadrature which is derived from the single-molecule annotation 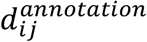 to get a combined distance: 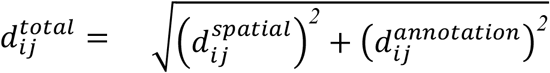. The annotation contribution can be either 0 within an annotation and infinity between annotations, automatically derived using distances between the expression profiles of the annotations, or manually specified. As the clustering works on affinities and not on distances, the distances have to be transformed to affinities, which is done by a Gaussian kernel with a subcellular distance scale.

As the number of molecules is so large, the clustering algorithm has to be fast and scalable, work entirely on sparse matrices, and cut the neighbourship graph at sensible places to generate reasonable cells-like objects. For this, a hierarchical clustering scheme was developed based on the spectral clustering implementation of scikit-learn: It can handle sparse matrices, can use algebraic multigrid solvers for speed, and, for only two clusters, it solves the normalized graph cut problem (*_spectral*.*py at 15a949460dbf19e5e196b8ef48f9712b72a3b3c3* · *Scikit-Learn/scikit-Learn*). The best cuts are found iteratively in top-down fashion (binary splitting the data at each iteration). To keep the dataset tractable for the initial cuts, the spatial structure of the data is used to generate super-nodes in the graph prior to clustering. When the clustering comes down into the size regime of a few single cells, several heuristics are employed to determine whether to accept a proposed cut based on the shape and size of the clusters, and on comparing the affinity loss of the proposed cut to the expected cost for cutting a homogeneous bulk graph of corresponding size and dimension. The final result is another column of annotation for the molecules containing unique object ids.

For visualization, these objects are filtered to include at least 20 molecules, and the associated expression profiles are normalized as in (Codeluppi et al. 2018), and embedded by a tSNE. To evaluate the self-consistency of annotation and segmentation, the silhouette score of the cell type annotations is evaluated on identically filtered and normalized profiles using the silhouette_score function from scikit-learn.

### Region definition

For spatially-convoluted expression data, such as in spatial transcriptomics methods including Slide-seq and Visium, clustering (of beads or spots) in expression space is less meaningful as individual cells are mixed. “Regions” which are defined both in position and expression or annotation space can be a meaningful alternative. TACCO implements a method to define such regions (**Supp. Fig. 3a**), consisting of two steps: (**1**) Construction of one *k*-nearest neighbors graph based on expression (or annotation) distances and another *k*-nearest neighbors graph based on physical distances; and (**2**) Combination of the two graphs as a weighted sum of the graphs’ adjacencies. Putting more weight on the position space adjacency gives contiguous regions in position space, while more weight on the expression adjacency leads to separated islands of similar expression being annotated with the same region and can connects regions across different spatial samples (*e*.*g*., Slide-seq pucks). To account for the missing links from the position graph between samples, the cross-sample adjacency is scaled up by a balancing weight factor. The combined adjacency matrix is then subjected to standard Leiden clustering(Traag et al. 2019) which assigns consistent region annotation across samples.

### Colocalization and neighborhood analysis

To score co-localization or co-occurrence of annotations (Palla, Giovanni, et al. 2021), TACCO calculates *p*(*anno*|*center*; *x*)/*p*(*anno*), the probability to find an annotation *anno* at a distance *x* from a center annotation *center* normalized by the probability to find *anno* regardless of a *center*. This is well-defined also for non-categorical annotations, which are commonplace for the compositional annotations created with TACCO and for pairs of unrelated annotations.

As the most time-consuming computations for co-occurrence are the same as for neighborhood enrichment analyses, TACCO also calculates neighborhood-enrichment z-scores (Keren et al. 2018) for a set of distance bins (as opposed to the set of direct neighbors on a graph), and again supports non-categorical annotation, pairs of unrelated annotations for *anno* and *center*, and multiple samples, while being competitive performance-wise (**Supp. Fig. 5**).

### Annotation coordinates

To analyze a given annotation (*e*.*g*. cell-type composition) with respect to its spatial distance from a reference annotation (*e*.*g*. histological annotation, **Supp. Fig. 3b,c**), TACCO implements an algorithm that determines a stable “annotation coordinate” in position space by regularizing the minimum distance by a critical neighborhood size determined at the weights level and afterwards correcting for the regularization bias. (This is required because for noisy spatial data and/or non-categorical reference annotations, simply taking the minimal distance between objects of certain annotations is unstable and/or not well defined.)

Specifically, TACCO first generates a matrix *n*_*A,x*_(*d*) of occurrence histograms *vs*. spatial distance *d* for every annotation category *A* and spatial position *x*, counting fractional annotations as fractional occurrence counts. The distance 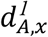 where its cumulative sum over distance *N*_*A,x*_(*d*) = Σ_*d*′_ *n*_A,x_(*d′*) will be over a certain threshold *N*_1_ is a robust measure for the radius of the sphere centered at *x* and containing a total of *N*_*1*_ occurrences of annotation *A*. Even for data homogeneously annotated with *A*, the minimal possible value of 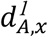 is however larger than *0* as the threshold *N*_*1*_ will in general be larger than *1* to have a stabilizing effect. In order to allow for a minimum value of *0* and obtain a measure for the distance from *x* to a significant amount of category 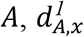, is corrected using a fictitious homogeneous annotation category *H*, the value of 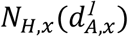 is found, followed by solving for the distance 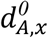 at which *N*_*1*_ less occurrences appeared: 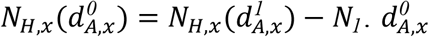 is now both consistent with the regular minimum distance for noise-free categorical annotations, stable against noise and well defined for compositional annotations.

.

### Enrichments

TACCO supports various approaches to visualize compositional differences in the spatial structure of a sample and estimate the statistical significance of these differences / enrichments (**Supp. Fig. 3d-g**). In particular, TACCO uses sample information to calculate enrichments not across observations (single cells/spatial beads, which are no independent observations and can lead to p-value inflation), but across multiple samples, which gives meaningful and reasonable p-values.

When there is just a single (or few) spatial sample(s) but with considerable size, different parts of that sample are treated as semi-independent biological replicates, by splitting the sample along a selection of coordinate axes to create a statistical ensemble for enrichment analysis. Increasing the number of splits to parts which cannot be regarded as independent replicates interpolates smoothly to where every observation is considered as independent.

### Datasets

#### Mouse colon scRNA-seq and *in-silico* spatial mixing

Mouse colon scRNA-seq data (Avraham-Davidi et al. 2022) in raw counts format after basic quality filtering, with the provided cell-type annotations, was used to simulate spatial mixing by sampling uniformly spatial coordinates for cells and for simulated beads on a square. To make boundary effects smaller, periodic boundary conditions were employed. On the resulting torus, the minimal Euclidean distance was taken between cells and beads. To factor in the “shape” of the beads, that distance was plugged into a kernel to compute the weights of contributions of each cell to a bead. From these weights, expression counts of cells, and cell-type annotation of cells, expression counts of beads, cell-type fractions, and count fractions belonging to each type were computed.

A Gaussian kernel was used to determine the weight of cell *c* in bead *b* as follows:

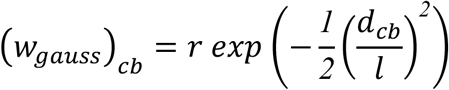

where *d*_*cb*_ is the Euclidean distance between the centers of cell *c* and bead *b*, 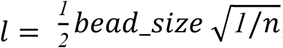 and *n* cells. To capture the higher sparsity of the spatial data, the weight was scaled by a capture rate parameter *r*, such that a cell contributes a fraction of *r* of its counts to a bead with distance *0. r* = *1*.*0* was used everywhere except where explicitly stated.

### Mouse colon scRNA-seq and Slide-seq data

scRNA-seq (as in the section above) and raw counts for Slide-seq data (Puck 2020-09-14_Puck_200701_21) of normal mouse colon were obtained from (Avraham-Davidi et al. 2022).

### Single-molecule osmFISH of the mouse cortex

osmFISH data was obtained from http://linnarssonlab.org/osmFISH/availability/ (Codeluppi et al. 2018). Reference expression profiles and their corresponding annotations were obtained from osmFISH-segmented cell profiles, provided in ‘osmFISH_SScortex_mouse_all_cells.loom’.

Raw mRNA locations were obtained from ‘mRNA_coords_raw_counting.hdf5’.

Preprocessing was performed using the procedure in https://github.com/HiDiHlabs/ssam_example/blob/master/osmFISH_SSp.ipynb. This includes: (1) Shifting and rescaling cell/RNA coordinates to match each other (with an RNA coordinate system); (2) Removing RNA molecules of bad quality; (3) Correcting incorrect gene names; (4) Removing molecules outside the intended imaged frame.

‘polyT_seg.pkl’, which contains a mapping of osmFISH’s segmented cell to molecule, was used to project osmFISH’s cell annotation onto individual molecules (approximating the ground truth). To visualize annotations at single-molecule level (**Fig. 2c**) a window of x range (1150,1400) and y range (1000,1600) was used.

For TACCO annotation, bin_size=10, n_shifts=3, bisections=4, bisection_divisor=3, platform_iterations=None were used.

For Baysor annotations, an adjacency graph was computed with build_molecule_graph, and cell types’ mean expression profiles (equivalent to the information used in TACCO) were provided for molecule annotation with cluster_molecules_on_mrf (do_maximize=false, max_iters=1000, n_iters_without_update=20).

### scRNA-seq simulations with scsim with enhanced dropout

Scsim(Kotliar et al. 2019) with its corresponding default parameters (changing deloc=5.0)^2^ was used to simulate scRNA-seq data. The dropout step described in Splatter(Zappia et al. 2017) was implemented in Python, by fitting a sigmoid curve through genes’ log mean count and their cell fraction with zero reads, where the sigmoid is characterized by shape and midpoint parameters. To enhance dropout, the sigmoid was shifted (decrease its midpoint) and the adjusted dropout probability was computed. This probability was then used to binomially sample the observed counts.

### scRNA-seq simulations with scsim with ambient RNA

To simulate expression profiles with ambient RNA, scsim(Kotliar 2019)‘s simulation of mean expression per cells and genes (termed “updatedmean” in scsim and denoted in Splatter(Zappia et al. 2017) as *λ*, a matrix of the means for each gene and each cell) was combined with CellBender(Fleming et al. 2019)‘s model of ambient RNA contamination and downstream sampling of counts.

Specifically, the mean expression of gene *g* and cell *n* (or here, the cell in drop *n*) was obtained from scsim as *λ*_*ng*_. For CellBender’s probabilistic model, the probability of having a cell in the drop *y*_*n*_ is set to 1 (because empty drops are not considered), and *ρ*_*n*_ is set to 0, because zero reads are exogenous to the drop as we account only for ambient RNA (pumped into the drop) and not for barcode swapping. Thus, the CellBender model simplifies to:

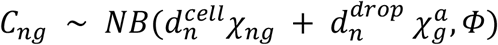

The two models were combined as follows:

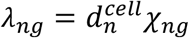is the mean expression (similar to CellBender, scsim also uses a log-normal distribution of cell size)

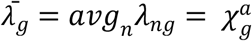is the mean true count of gene *g*, that is, the ambient contribution of the gene

Given the fraction of ambient RNA, *f*^*drop*^, and the library size, 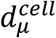, and scale, 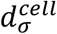, used for sampling the cell size, 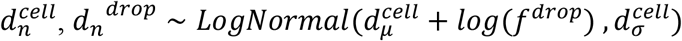 is sampled (that is, the same variance is employed as used for sampling the cell content and the mean is set to 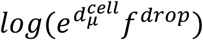).

*Ф*is sampled as defined in CellBender

Thus, we sample:

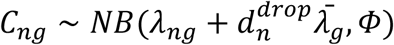

For comparability, the reference is generated in the same way (using negative binomial sampling instead of scsim’s Poisson sampling), but without adding ambient RNA.

### scRNA-seq of clone-labeled, differentiating hematopoiesis cells

Normalized counts (‘stateFate_inVitro_normed_counts.mtx’, together with the corresponding gene names and cell metadata) and cells’ clone matrix (‘stateFate_inVitro_clone_matrix.mtx’) were obtained from the github repository https://github.com/AllonKleinLab/paper-data/blob/master/ (Weinreb et al. 2020). Cells were filtered to those belonging to clones with cells in day 2 and in days 4 or 6. The fate bias for each cell in day 2, was computed from the fates of its clone in days 4 and 6 (normalized to 1). Differentiated cells captured in days 4 and 6 were then used as a reference for annotating cells of differentiated fate from day 2.

### TACCO configuration

For all datasets, TACCO was applied with basic platform normalization, entropy regularization parameter epsilon 0.005, marginal relaxation parameter lambda of 0.1, 4 iterations of boosting with a divisor of 3 and using 10 representing means for each type. The only exception is **Fig. 4** where in addition to TACCO with these parameters, TACCO was also applied without boosting, using only the maximum annotation per observation as categorical annotation, and all other parameters identical as an optimized categorical annotation method “TACCO (cat)”.

## Code Availability

TACCO is available as the open-source python package tacco, with source code freely available at https://github.com/simonwm/tacco and corresponding documentation at https://simonwm.github.io/tacco/.

## Acknowledgements

We thank L. Gaffney for help with figure preparation. S.M. was supported by a DFG research fellowship (MA 9108/1-1), J.K. was supported by a HFSP long term fellowship (LT000452/2019-L), A.R. was a Howard Hughes Medical Institute (HHMI) Investigator when conducting this work. The work was supported by the Klarman Cell Observatory, a CEGS grant (5RM1HG006193-09) from the NHGRI, the NIH/NIAID (grants 1U24 CA180922, 1U19 MH114821, 1RC2 DK114784), the MIT Ludwig Center, the Manton Family Foundation, and HHMI (A.R.); Azrieli Foundation Early Career Faculty Fellowship, and an ISF Research Grant (1079/21) (M.N.), and the Center for Interdisciplinary Data Science Research at the Hebrew University of Jerusalem (N.M. and M.N.).

## Author contributions

N.M., I.A-D., M.N. and A.R. conceived the study. S.M., N.M., J.K. and M.N. developed the algorithms and framework, with input from I.A-D. and guidance from A.R.; S.M. and N.M. implemented the software and performed the analyses and benchmarks; I.A.-D., E.M. and F.C. provided mouse colon Slide-Seq data; I.A.-D., O.R.-R. and A.R. provided mouse colon single-cell RNA-seq data; I.A.-D., J.K. and A.R. assessed and interpreted biological applications; J.K., A.R. and M.N. supervised the research; S.M., N.M., I.A.-D., J.K., A.R. and M.N. wrote the manuscript.

## Supplementary Figures

**Supp. Figure 1.**
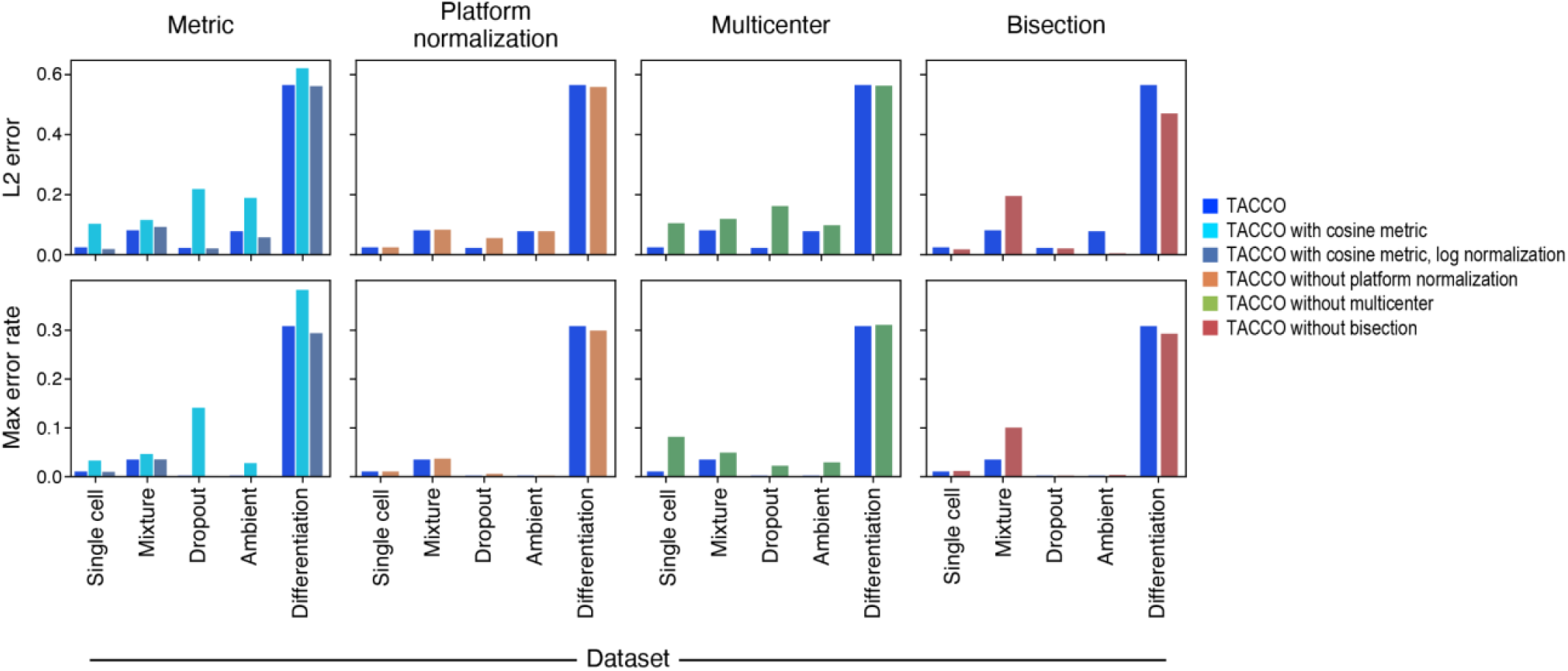
Effect of choice of cost matrix metric in OT and several booster options (platform normalization, multicenter, bisection) on annotation quality. L2 error (y axis, top; relevant for mixture annotations) and fraction of objects where the maximum annotation disagrees with the reference (y axis, bottom; relevant for classification problems) for different methods (colors) on different use cases (x axis). Baseline TACCO used the Bhattacharyya metric, platform normalization, multicenter, and bisection. Single cell: annotation of mouse colon scRNA-seq with itself as reference; Mixture: Gaussian mixtures of mouse colon scRNA-seq (**Fig. 2a**); Dropout: simulated Dropout dataset (**Fig. 3a**); Ambient: simulated ambient dataset (**Fig. 3b**), Differentiation: hematopoiesis scRNA-seq (**Fig. 3c**).

**Supp. Figure 2.**
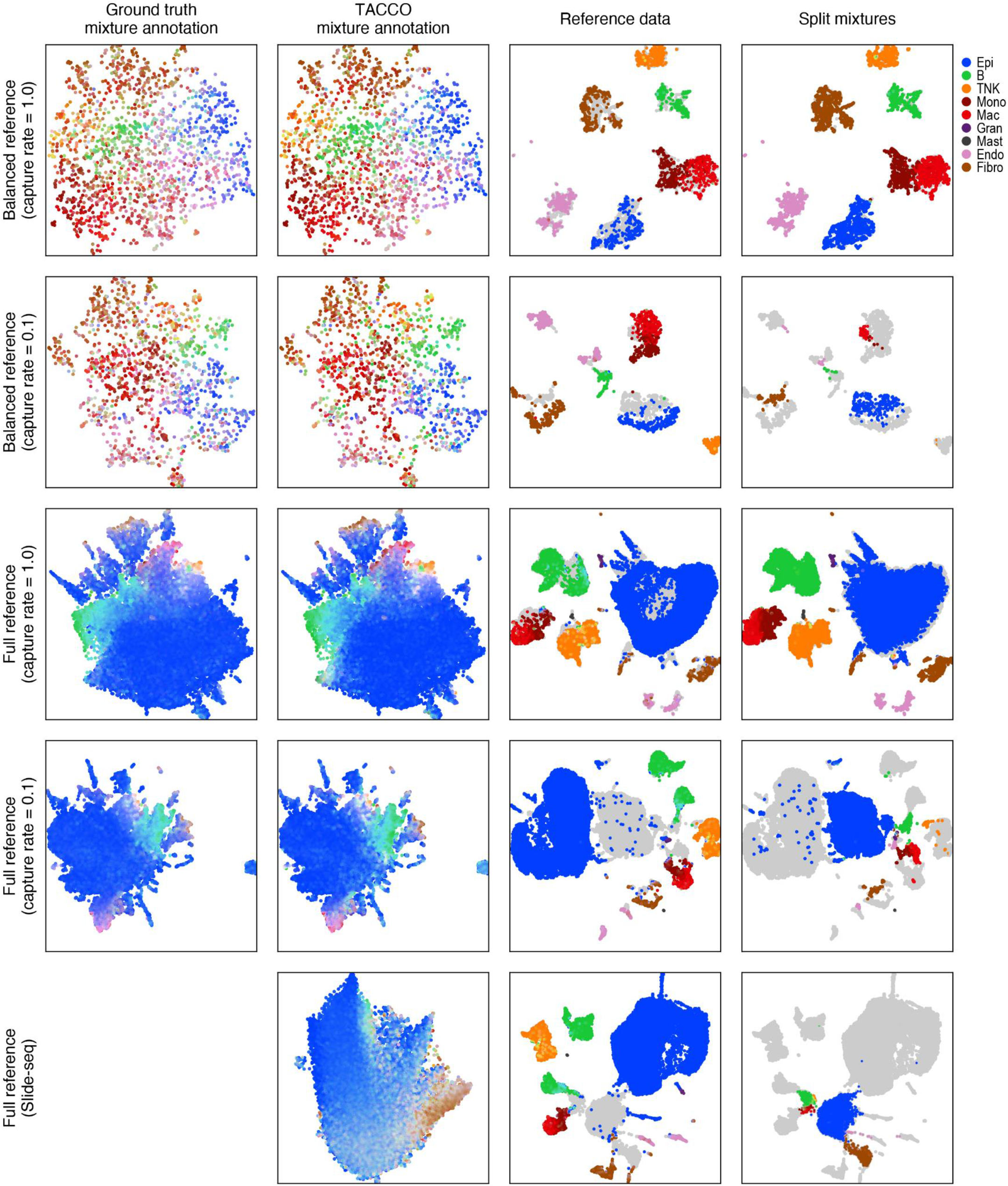
TACCO splitting of mixed expression profiles into pure constituents. UMAP embeddings of in-silico mixed mouse colon scRNA-seq profiles (top four rows: real or balanced type composition and with and without downscaling the count-yield per cell by a factor of 10) or Slide-seq beads (bottom row), based on ground truth (far left, where available), TACCO annotation (second left), reference data (second right) and mixed expression data split using TACCOs annotation as input embedded in a joint UMAP of the concatenated reference-split dataset (rightmost). The in silico mix uses a beadsize of 1.0. For the joint embedding data was filtered to include only observations with at least 30 counts in genes which were annotated highly variable in the reference dataset, except for capture_rate=1.0 where a threshold of 100 is used.

**Supp. Figure 3.**
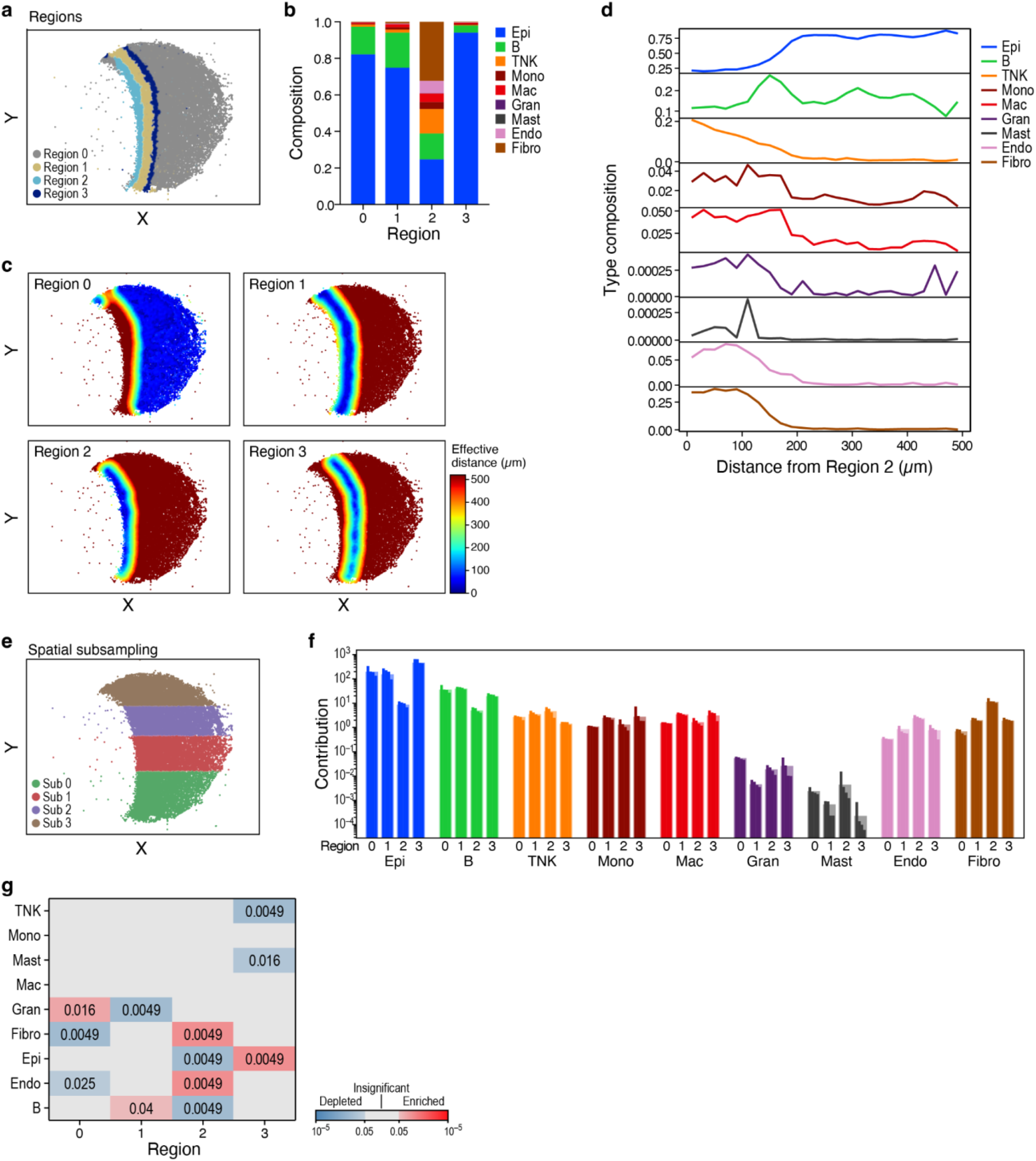
Example workflow and visualizations for spatial data analysis in TACCO for the mouse colon dataset from the “Spatial mixture” task (Fig. 2b): (a) Slide-seq puck with beads colored by regions defined by joint expression space and position space clustering. (b) Fraction of cells (y axis) of each type (color) in each region (x axis). (c) Slide-seq puck colored by the effective distance of each bead from each region. (d) Density (y axis) of cell type annotations at different effective distance from region 2 (x axis). (e) Slide-seq puck colored by spatial split of the puck along a selected coordinate axis into multiple biological replicates for statistical enrichment analysis. (f) Geometric mean normalized cell type fractions (y axis) for each cell type (x axis) from each region (1-4) and subregion (per spatial split as in e; individual bars). (g) Enrichment p-values (colors) of the normalized cell fractions in (f) (Benjamini-Hochberg-corrected one-sided Mann-Whitney-U test across spatial splits).

**Supp. Figure 4.**
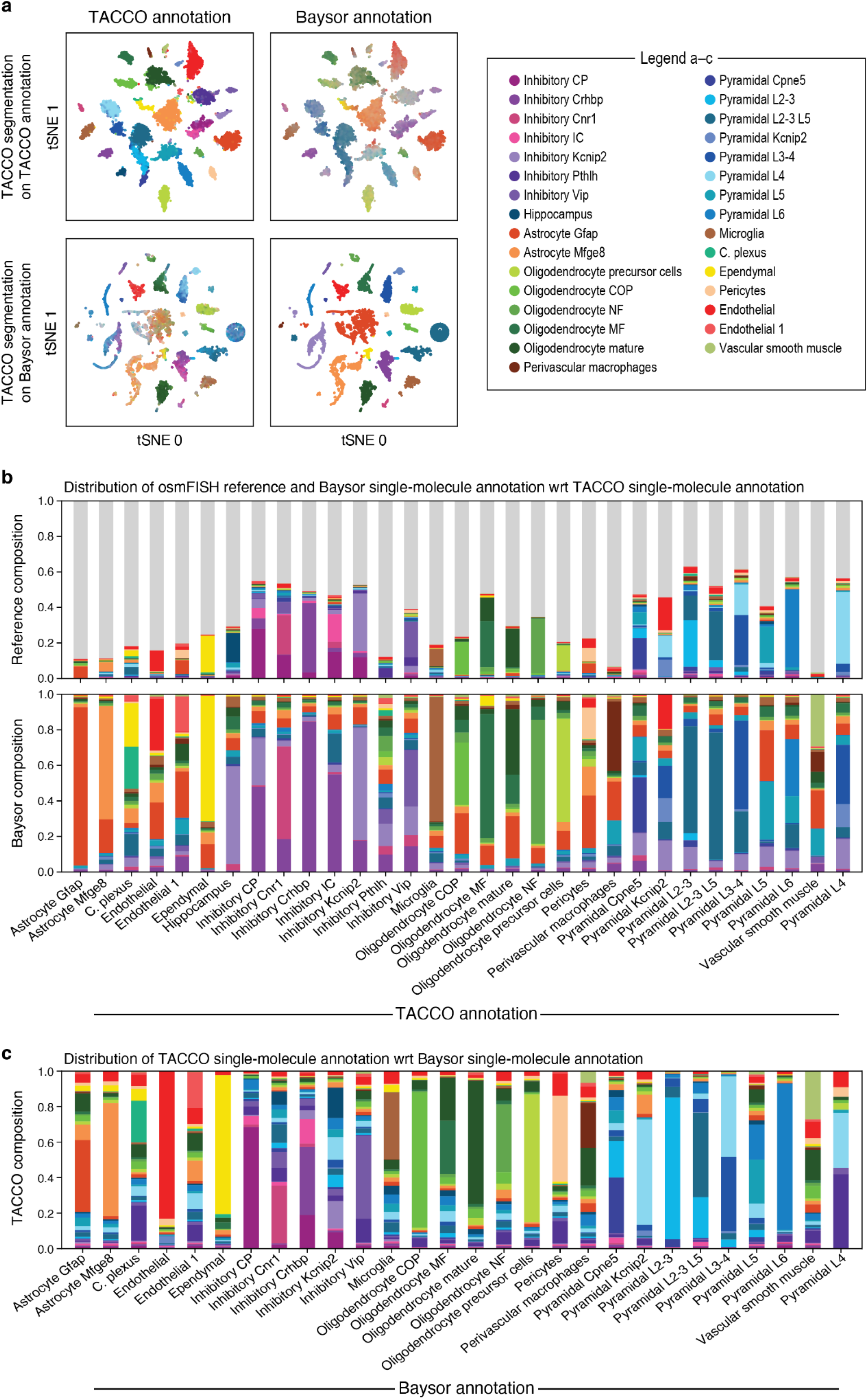
Comparisons of single-molecule annotations for osmFISH(Codeluppi et al. 2018). (a) tSNE embeddings of the expression profiles for objects (dots) resulting from TACCO segmentation using either TACCO annotation (top row) or Baysor single molecule annotations as input (bottom row) with objects colored by the aggregated single-molecule annotation from either TACCO (left) or Baysor (right); (b) distribution of reference annotation and Baysor single-molecule annotation over the TACCO single-molecule annotations; (c) distribution of TACCO single-molecule annotation over the Baysor single-molecule annotations.

**Supp. Figure 5.**
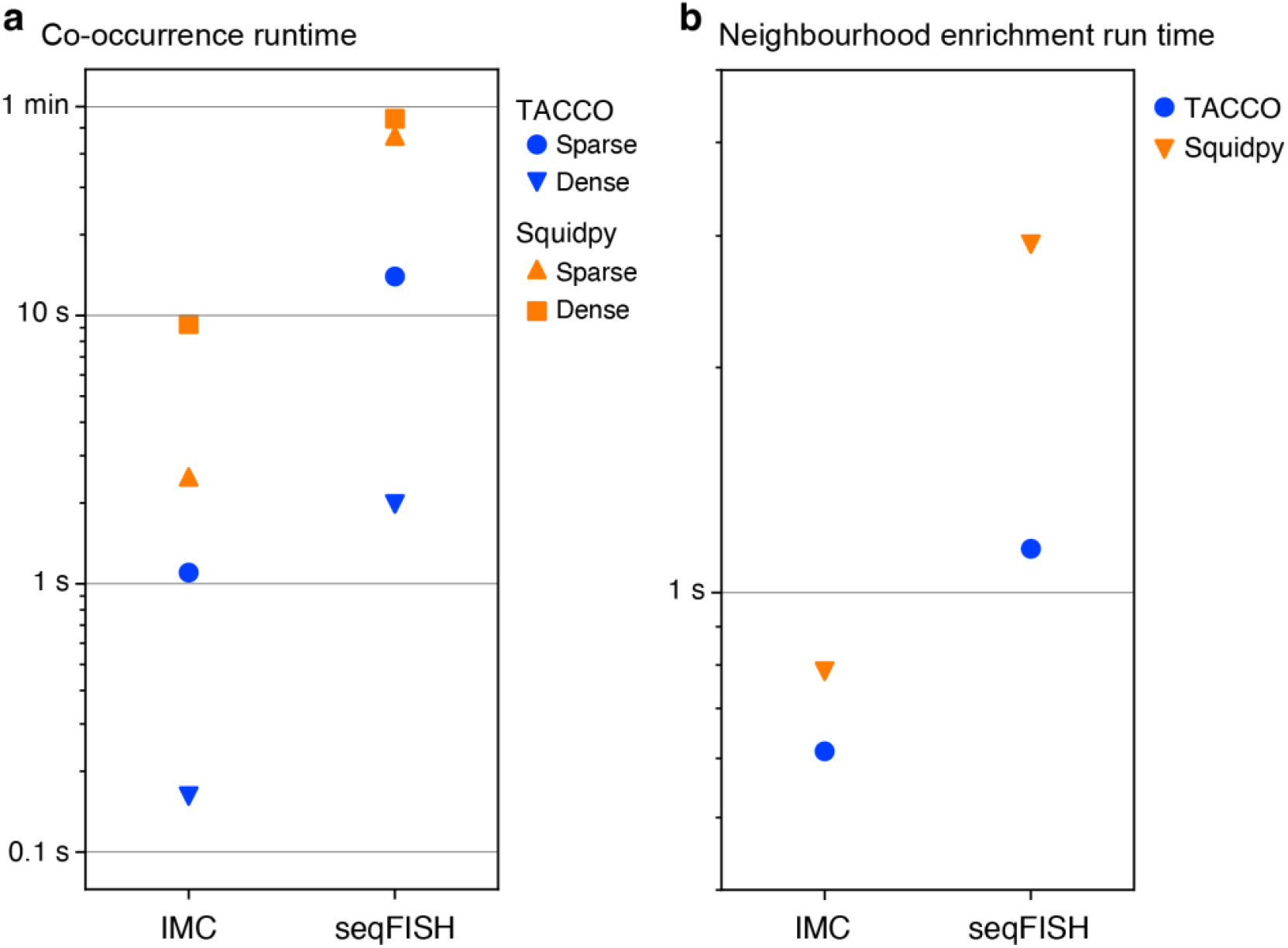
Runtime comparison of TACCO and Squidpy(Palla et al. 2021). Run time (y axis) co-occurrence (a) and neighbourhood enrichment (b) analyses for TACCO (blue) and Squidpy (orange). Datasets “imc” and “seqfish” are example datasets provided with Squidpy.

## Footnotes

1 Setting n(tb) to p(tb) one can reproduce the result from Cable et al. by determining n(gb) from the t-marginal, ignoring the g-marginal *i*.*e*. the annotation result.

2 Taken from https://codeocean.com/capsule/6314882/tree/v1, code/analysis/Part1_Simulations/Step1_Simulate.ipynb

